# Volumetric and Diffusion Tensor Imaging biomarkers indicating long lasting post-concussion abnormalities in a youth pig model of mild Traumatic Brain Injury

**DOI:** 10.1101/2024.11.12.623259

**Authors:** Islam Sanjida, Netzley Alesa, Li Chenyang, Zhang Jiangyang, Dávila-Montero Bianca, Vazquez Ana, Subbaiah Shaun, Meoded Avner, Munoz Kirk, Colbath Aimee, Huang Jie, Mejia-Alvarez Ricardo, Manfredi Jane, Pelled Galit

## Abstract

Mild Traumatic Brain Injury (mTBI) caused by sports-related incidents in children and youth can lead to prolonged cognitive impairments, underscoring the importance of improved diagnosis and comprehension of its enduring impacts on neuropathology. A pig model was chosen for its similarities to the human brain in terms of gyrencephalic structure, size, and regional proportions, and a closed-head mTBI was induced in adolescent pigs. In this study, 12 (n=4 male and n=8 female) 16-weeks old Yucatan pigs were tested; n=6 received mTBI and n=6 received a Sham procedure. This study utilized T1-weighted imaging to assess volumetric alterations in different regions of the brain and diffusion tensor imaging (DTI) to examine microstructural damage in white matter. The pigs were imaged at one and three months post-injury. Our volumetric analysis of key white and gray matter regions showed significant longitudinal changes in pigs with mTBI compared to sham controls. The observed volume increases may be attributed to swelling, neuroinflammation, or hyperactivity. Fractional anisotropy (FA) values derived from DTI images demonstrated an increase in corpus callosum from 1 month to 3 months only in mTBI pigs. Additionally, comparisons of the left and right internal capsules revealed a decrease in FA in the right internal capsule for mTBI pigs, likely due to the impact being slightly localized to the right side of the brain, which may indicate demyelination. Thus, the injury has disrupted the maturation of white and gray matter of the developing brain. This signifies the need for longitudinal investigations after mTBI to comprehensively assess its long-term effects and contribute to the clinical management of concussion in youth.

## Introduction

Traumatic brain injury (TBI) constitutes a significant global public health challenge, posing severe and widespread implications for neurological health and healthcare systems worldwide. TBI often results from exposure to blasts, falls, motor vehicle accidents, sports-related injuries, or combat situations. Globally, TBI is a significant contributor to the loss of disability-adjusted life years. TBI affects millions, including children, leading to significant cognitive deficits and fatality. Annually, 475,000 children in the U.S. experience traumatic brain injuries, leading to 37,000 hospitalizations and 2,685 deaths, with the highest emergency room visits and death rates in children under 4 years.^1^

Mild TBI (mTBI) is synonymous with concussion^2^ and is most prevalent in children and youth due to their active participation in sports.^3^ Symptoms of mTBI can include nausea, headaches, and possible loss of consciousness for several minutes. Since these symptoms often resolve rapidly, mTBI diagnosis remains challenging. A substantial number of children (25.3% in the mTBI group^4^) experience persistent post-concussion syndrome (PCS).^5,6^ PCS symptoms which include headaches, dizziness, fatigue, anxiety, depression, difficulty concentrating, and memory problems can last for weeks, months, or even years after a concussion.^7^ The severity of PCS symptoms was correlated with objectively measurable deficits in neurocognitive function, indicating that mTBI can have long-term consequences and that PCS is not merely a transient condition.^8^ The study found that a significant proportion of TBI patients reported persistent symptoms one month after the injury and an even greater number of patients reported persistent symptoms a year later.^9^ Even among mTBI patients, 44% had three or more symptoms one year after the injury, indicating higher rates of persistent symptoms than commonly believed.^10^ Repeated mTBIs sustained in contact sports are strongly linked to the onset of chronic traumatic encephalopathy, a progressive neurodegenerative disorder marked by the pathological accumulation of tau protein within the brain.^11,12^

Although standardized evaluation tools are helpful for medical management of mTBI, the progression and timeline of recovery from mTBI or Sports Related Concussion (SRC) remains unclear.^13,14^ Consequently, non-invasive techniques such as MRI and CT scans have become standard tools for evaluating clinical outcomes and identifying biomarkers.^15^ mTBI is a multifaceted condition that significantly impacts brain function. Although the precise mechanisms are still under investigation, the pathophysiology of mTBI likely involves a combination of mechanical forces, biochemical changes, and functional consequences.^16^ Mild injuries can lead to axonal damage, which may later result in axonal separation. Mechanical forces during trauma that damage the axonal cytoskeleton. Initially, this disruption impairs axonal transport, leading to axonal swelling^17^ and ultimately causing Wallerian degeneration, which can progress over months to years, post-injury. Demyelination is also an important feature of axonal damage which refers to the gradual death and damage of myelin sheaths of axons disrupted from the trauma or the external force in mTBI.

Anatomical MRI and diffusion tensor imaging (DTI)^18^ provide means to assess white matter pathology based on regional volumes and microstructural integrity.^19-21^ The primary DTI parameters include fractional anisotropy (FA), mean diffusivity (MD), axial and radial diffusivity (AD and RD),^22^ which provide valuable but indirect measurements of white matter integrity. For example, previous studies suggest that FA values increase with axonal density and myelination. MD values decrease under axonal swelling and reduce extracellular water.^23^ Changes in AD and RD have been associated with axonal and myelin injuries.^24,25^

Numerous studies have explored DTI analysis following mTBI, employing methods such as region-of-interest (ROI) and whole-brain or voxel-based approaches.^26^ While ROI analysis is widely used and advantageous, it has limitations for TBI research due to the diffuse nature of axonal injury. However, abnormalities in diffusion and diffusion anisotropy in mTBI are frequently too minor to be identified through whole-brain histogram analysis.^27^

To unravel the complexities of mTBI researchers turn to animal models which enable a reduction in heterogeneity, identification of performance of longitudinal studies to assess test interventions, thus facilitating understanding of pathophysiological mechanisms which can and identify therapeutic targets.^28-30^ Rodent models of injury including the weight drop,^31-33^ fluid percussion injury (FPI),^34^ controlled cortical impact (CCI),^21,35-38^ closed head impact model of engineered rotational acceleration (CHIMERA),^39^ and blast TBI,^40^ have significantly added to the understanding of cellular and systemic mechanisms associated with injury. The pig became an established experimental model for head injury,^41-48^ recapitulating many of the important behavioral and pathophysiological changes experienced by humans and allowing for longitudinal neuroimaging with neurocognitive and physiological screening. Previously we characterized the cognitive and behavioral performance of Yucatan minipigs throughout their development which is essential to serve as a reference point for behavioral function.^49^ Here, we developed a closed head mTBI model in adolescent minipigs. An electromagnetically driven impactor delivered an injury with a clinically relevant magnitude. T1-based MRI contrast and DTI were used to characterize post-concussive gray and white matter integrity in the months after the mTBI. The results demonstrate DTI abnormalities in pigs that underwent mTBI even months after the injury, which may be contributing to long-lasting functional deficits. This work could aid in developing better diagnostic tools and treatments for mTBI in youth.

## Methods

### mTBI model

All procedures were approved by the Institutional Animal Care and Use Committee at Michigan State University. 16-week-old male (n=4) and female (n=8) Yucatan miniature pigs were used in this study. Pigs were housed in an enriched environment, fed nutritionally complete feed twice per day, with unrestricted access to water, while on a 12-hour (7:00-19:00) light cycle. Each cohort included four individuals, two who underwent the mTBI procedure and two who underwent the Sham procedure. Each cohort was housed together throughout the study. Prior to surgery, pigs were administered midazolam (0.2-0.4 mg/kg) and butorphanol (0.2-0.4 mg/kg) intramuscularly. Pigs were mask induced, intubated and inhalant anesthesia was used to maintain a surgical plane of general anesthesia. End-tidal CO_2_, hemoglobin oxygen saturation, blood pressure, heart rate, respiratory rate, and core body temperature were monitored throughout the duration of the surgical procedure and any abnormalities were corrected. Once a surgical plane of anesthesia was achieved, a skin and periosteal flap was created over the right coronal and sagittal suture. A custom designed sterile 3D printed disk (made of a digital blend of Vero Clear and Agilus30 (Stratasys)) was placed on the skull to protect the skull from fracture upon impact. Following impact, the periosteum and skin were closed in separate layers. Inhalant anesthesia was discontinued, and the pigs were extubated after they showed signs of recovering from anesthesia. Sham-control pigs were anesthetized and intubated for the same duration of the injury protocol and underwent the same skin and periosteal incision and closure and received the same perioperative and operative medications. Pigs were pain scored after their surgical procedure to determine if additional analgesics were needed.

We developed an electromagnetically driven impactor to deliver highly controlled, reproducible impacts to the pigs’ heads utilizing a linear motor (PS01-37X120F-HP-C, LinMot USA Inc., WI, USA). This device enables precise adjustment of injury severity, as quantified by the Severity Index (SI), to simulate and study the mechanical forces involved in mTBI. The impactor (12.55 mm in diameter and 600 g in mass) was placed on the right side of the exposed skull at Bregma 0. Ultra-high-speed camera (Phantom V2512 Series) was used to record the impact kinematics and subsequent subject’s movement at 20,000 frames per second with a spatial resolution of 1024 × 1024 pixels. An in-house Matlab (R2018b) code was built to track specific features of interest in the frame utilizing a region growing algorithm approach. In order to visualize the linear momentum effects upon impact, the impactor tip and subject’s eye were utilized as features of interest.

### Data Acquisition

MRI was performed at 1 and 3 months after mTBI. Pigs underwent anesthesia as described above, and while in the MRI they were maintained on a continuous intravenous infusion of propofol and inhaled oxygen. All data were acquired on a Siemens 1.5T system. The protocol included axial T2-weighted fluid-attenuated inversion recovery (FLAIR), DTI, axial gradient-echo (GRE), and 3D axial T1-weighted scans. The 3D-T1 scan was acquired using the following parameters: TE/TR=3.02/2400 ms; FOV 187×250 mm^2^; matrix 192×256; slice thickness 1 mm; voxel size 0.9766×0.9766×1 mm^3^; acquisition time 15.4 min. The DTI sequence was acquired in the axial plane, using 64 directions and one b0 using the following parameters: TE/TR=127/7900 ms, NEX=1, FOV=230 mm, matrix 114×114, slice thickness 2 mm, voxel size=2 mm^3^, 40 slices, b=1000 s/mm^2^, and acquisition time of 8.72 min. Two DTI scans were acquired: one with phase encoding from anterior to posterior and the other from posterior to anterior.

### MRI Analysis

Manual Brain Skull stripping: The FSL package (Analysis Group, FMRIB, Oxford, UK) was used to create brain masks and registration. Brain extractions were performed by a single evaluator to minimize variability, without prior image intensity normalization before manual extraction. Representative brain masks are shown in **Figure 1**.

**Figure 1.**
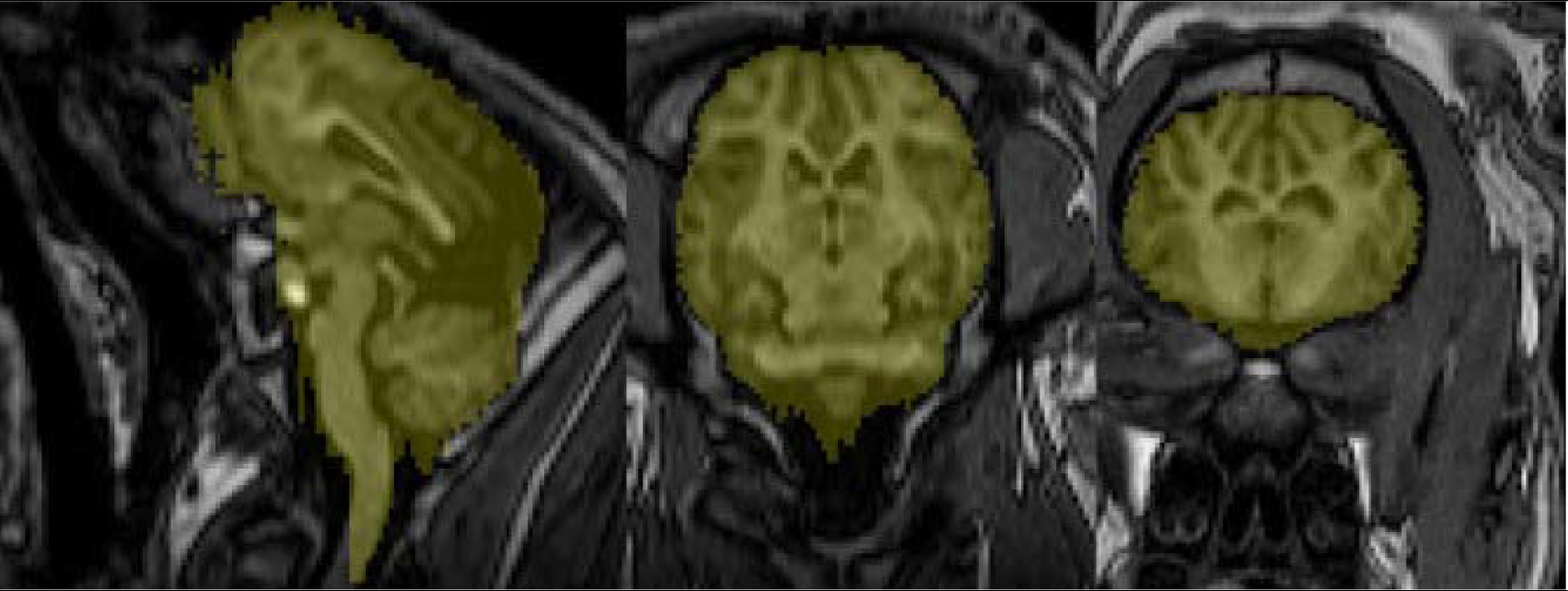
T1-Weighted MRI scans. T1-weighted MRI scans of pig brains illustrating manual brain skull stripping across different planes (from left to right, Sagittal, Coronal and Axial plane, respectively). The planes show the brain with the manually drawn mask overlay used for skull stripping. The mask highlights the brain’s boundaries, excluding the surrounding skull and non-brain tissues.

### Image Processing and Analysis

Initially, all diffusion images underwent preprocessing using FSL and MRtrix. This preprocessing encompassed several sequential steps: denoising to reduce noise from diffusion-weighted images, followed by Gibbs ringing removal. Subsequently, motion and distortion and eddy current corrections were performed, brain masks were generated and applied to extract brain data from the preprocessed images. Finally, the diffusion tensor was computed, along with metrics including FA, MD, RD, and AD. Our methodology utilizes an automated, atlas-driven approach, which minimizes the inconsistencies linked with manual region selection and improves reliability. We used a pig brain atlas^50^ to examine changes in regional volume and DTI metrics in this study. The atlas is based on group-averaged MR images of the pig brain and provides a parcellation map for major gray and matter structures. Registration of the atlas to the T1 scans were performed using the “antsRegistration” program from the Advanced Normalization Tools (ANTs). The ROIs in the atlas were applied to co-registered T1 and DTI data using the “antsApplyTransforms” program to measure structural volumes and mean FA, MD, RD, and AD values within each ROI.

### Histopathology

At the endpoint, the brains were removed and placed overnight in 4%PFA. Ten sections of 4 µm brain sections were obtained from the frontal cortex every 500 mm, and 10 sections were obtained every 500 mm around Bregma 0, where the insult occurred. Staining for Luxol fast blue (LFB) was performed. Olympus FluoView 1000 LSM was used to image slides. Three to five images were acquired with a magnification of 20X. The percentage of pixels indicating cellular uptake of stain was calculated using a custom macro with ImageJ (NIH) for each image.

### Statistical Analysis

Analyses of DTI parameters, along with variations in white matter and gray matter volumes, were executed using statistical modules available in GraphPad Prism. We hypothesized that individuals with mTBI demonstrate white matter tract damage, evidenced by changes in FA and volume, relative to the sham group at the 1-month and 3-month time points. To test this hypothesis, cross-sectional comparisons were conducted with two sample t-tests for each respective time point. Furthermore, to explore longitudinal changes within each cohort, we employed paired comparisons or paired t-tests for both the mTBI and sham groups. Moreover, to control for Type I error, p-values were adjusted using the Bonferroni correction method, accounting for the number of tests conducted in the multiple comparisons of the Volumetric Analysis. To investigate the impact of gender, we performed a mixed model analysis. This method enables us to consider fixed effects, such as gender, group, and time point, while also accounting for subject variation as random effects.

## Results

### Kinematics of Injury

Full displacement of the subject and impactor was recorded to characterize the impact kinematics and mechanical response upon impact. Tracking the impactor showed that it reached a peak speed of 3.7 m/s with an average 30g acceleration. Using the pig’s eye as a marker, its displacement and velocity were derived to obtain an experienced maximum acceleration of approximately 15g. Experimental data demonstrating the eye’s displacement, velocity, and acceleration due to the impact of three trials is illustrated in **Figure 2**.

**Figure 2.**
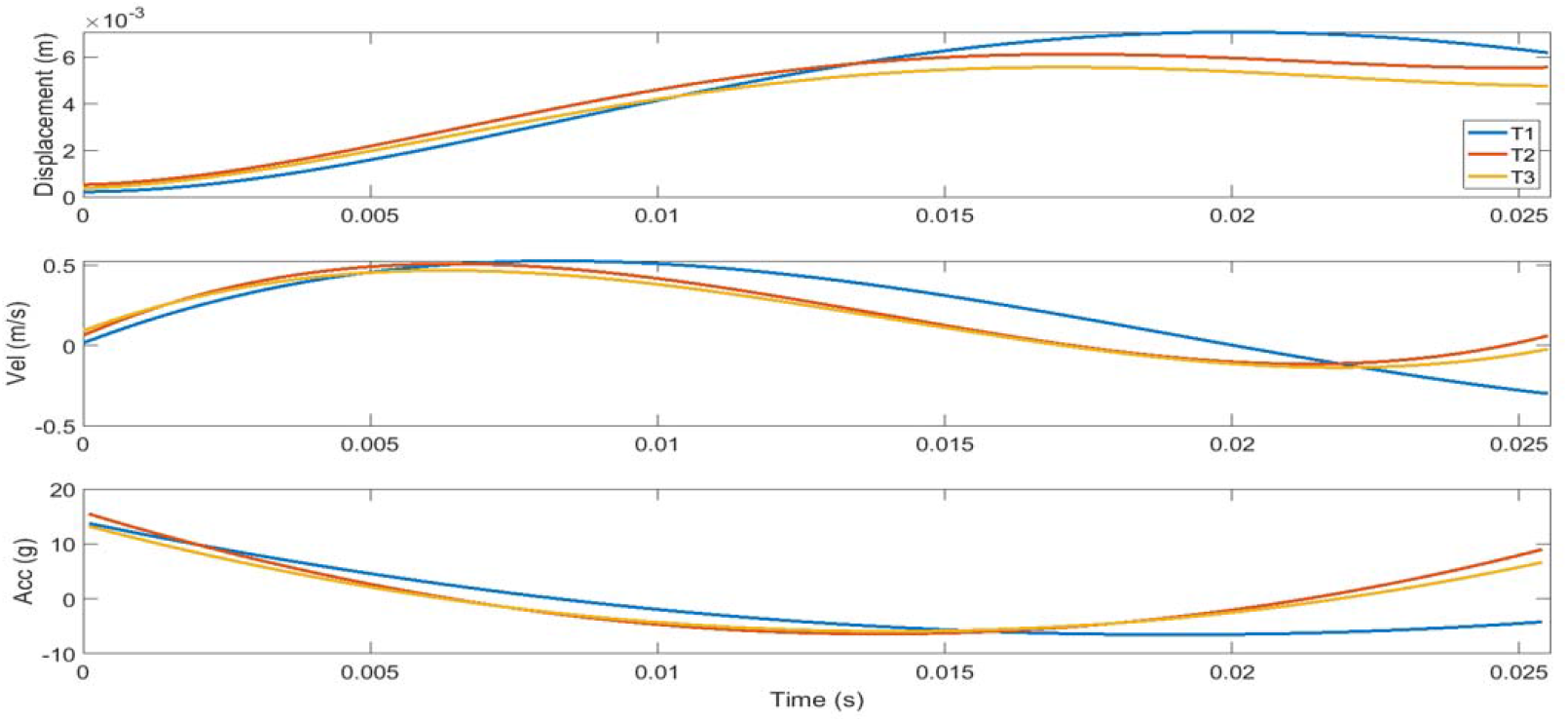
Kinematics of head injury. Kinematics of injury for head’s rigid motion using the eye as tracking marker. Displacements were calculated with a MATLAB custom-built code for image processing, applying a polynomial fit and then derived for velocity and acceleration.

### Volumetric analysis

The volume of the brain region can be an important indicator of abnormal function associated with brain trauma. For the volumetric analysis, both white matter and gray matter regions were examined. **Table 1** shows the results of the volumetric analysis.

**Table 1:** **Volume Changes in mTBI and Sham pigs across brain regions**: 1-month and 3-month analysis with percentage change and Paired t-Test results (*, Significant at p<0.05; and **, Significant at p<0.003 (0.05/18), where 18 is the number of tests.)

| Region of Interest | mTBI |  |  |  | Sham |  |  |  |
| --- | --- | --- | --- | --- | --- | --- | --- | --- |
| | 1-month<br>(mean ±SD)<br>mm<sup>3</sup> | 3-month<br>(mean ±SD)<br>mm<sup>3</sup> | % of<br>change | P value | 1-month<br>(mean ±SD)<br>mm<sup>3</sup> | 3-month<br>(mean ±SD)<br>mm<sup>3</sup> | % of change | P value |
| Corpus Callosum | 1104±102 | 1228±91.1 | 11.23188<br>406 | 0.00124** | 1059±158.0 | 1182±169.9 | 11.61473088 | 0.05899 |
| Left Internal Capsule | 1672±152 | 1822±153.6 | 8.971291<br>866 | 0.00612* | 1657±168.5 | 1839±236.1 | 10.98370549 | 0.049313* |
| Right Internal Capsule | 1728±139 | 1904±197.1 | 10.18518<br>519 | 0.00713* | 1698±188.2 | 1882±202.2 | 10.83627797 | 0.039794* |
| Left Cortex | 37140±27 | 39411±2819.8 | 6.114701<br>131 | 0.00803* | 37071±4532.6 | 40312±6125.6 | 8.74268296 | 0.088732 |
| Right Cortex | 36638±25 | 39191±3069.9 | 6.968175<br>119 | 0.00362* | 36430±3908.1 | 39576±6042.1 | 8.635739775 | 0.084723 |
| Left Hippocampus | 1122±181 | 1226±180.4 | 9.269162<br>21 | 0.00546* | 1112±122.4 | 1158±163.9 | 4.136690647 | 0.575766 |
| Right Hippocampus | 1131±148 | 1232±134.9 | 8.930150<br>309 | 0.00163* | 1121±149.3 | 1174±239.5 | 4.727921499 | 0.507322 |
| Cerebellum | 15331±101 | 15934±903.7 | 3.933207<br>227 | 0.098859 | 15287±2380.5 | 14539±2495.1 | 4.893046379 | 0.462634 |
| Thalamus | 2843±166.6 | 2994±126.8 | 5.311290<br>89 | 0.010258* | 2856±269.4 | 2972±314.7 | 4.06162465 | 0.449299 |

The volume of the corpus callosum in mTBI pigs was found to significantly increase between 1-month to 3-months after injury, which is consistent with the FA analysis. The data in the table indicate that all white matter regions, including the Left and Right Internal Capsule, exhibited volume increases over time in both mTBI and Sham groups. However, the increase was more pronounced in the mTBI group.

In addition, the majority of gray matter regions in the Left and Right Cortex, Left and Right Hippocampus, and Thalamus exhibited a notable increase in volume in the mTBI groups. In contrast, the cerebellum did not demonstrate any volumetric changes in either the mTBI or Sham groups (**Figure 3**).

**Figure 3.**
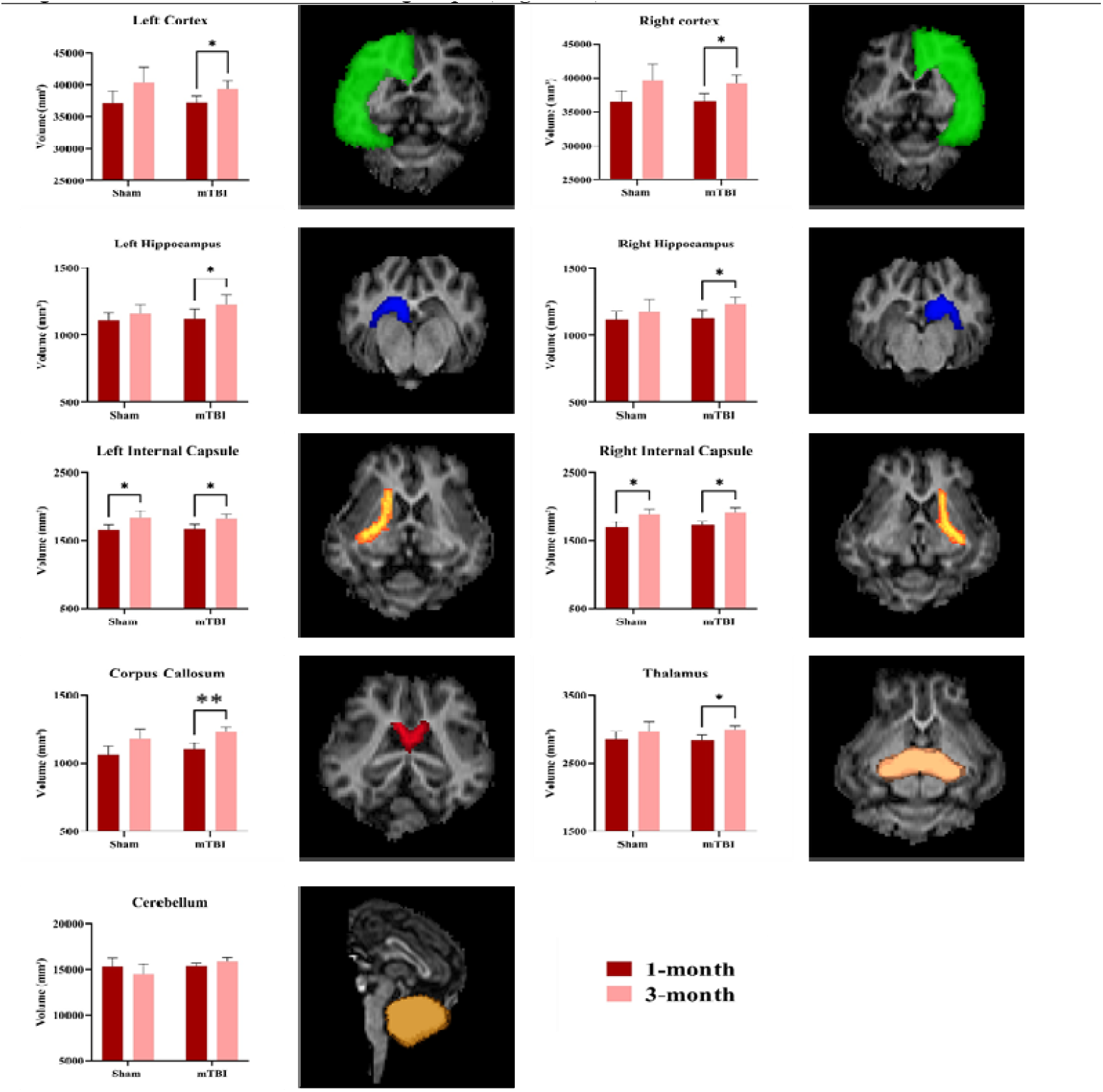
Volumetric analysis. Displays representative maps of the white and gray matter regions alongsid their corresponding group analysis results (mTBI, n=6; Sham, n=6; avg±SEM; Student t-test, *, p<0.05; **, Significant at p<0.003).

We conducted an additional analysis to compare volume changes between the left and right internal capsules in pigs with mTBI compared to sham. Consistent with the DTI results, volume alterations were observed solely in the mTBI pigs at both 1-month and 3-months. Notably, the volume of the right internal capsule seemed to increase more than that of the left as shown in **Figure 4**.

**Figure 4.**
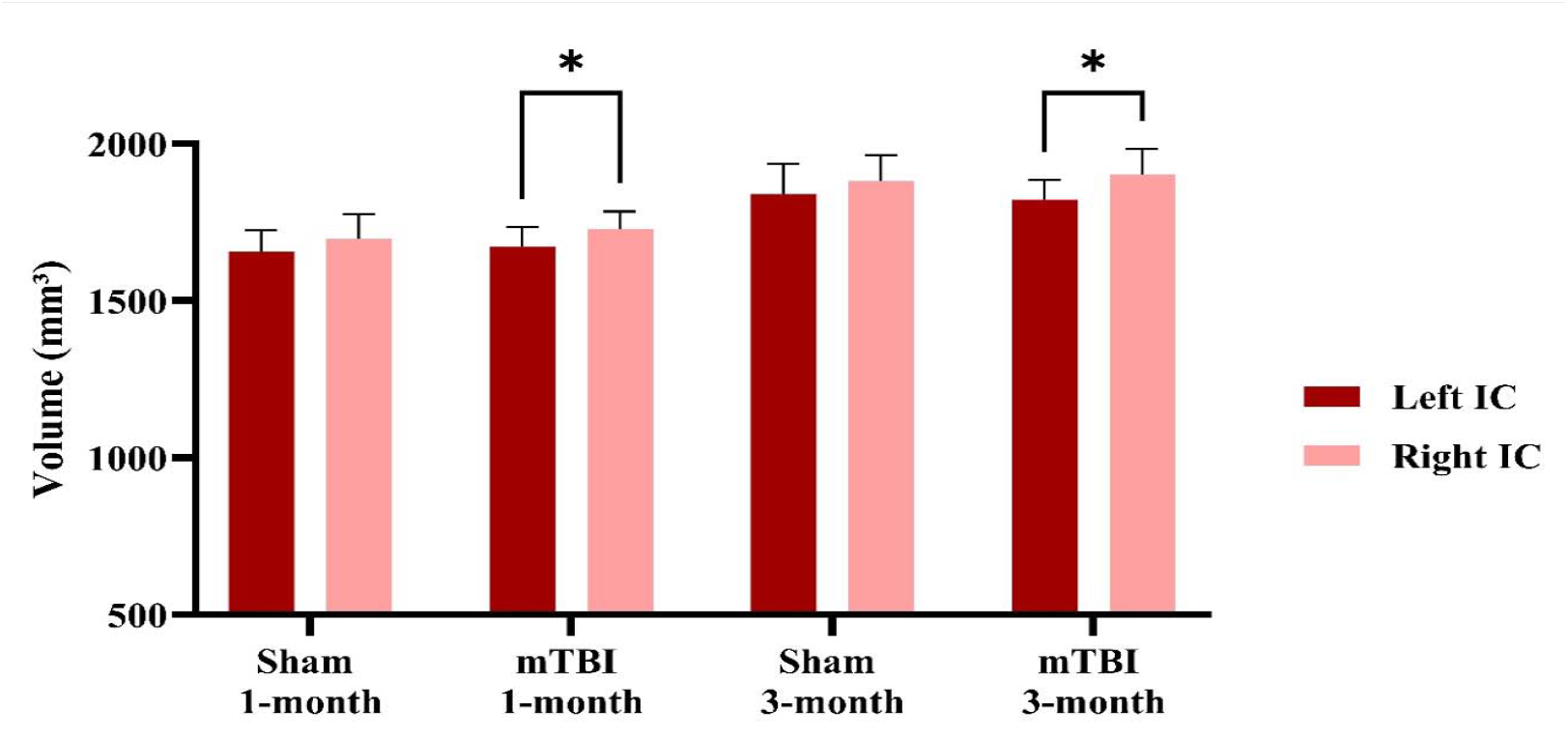
Volumetric differences in Left and Right Internal Capsule post-mTBI. This volumetric analysis of the Left and Right Internal Capsule. The results demonstrate that in the mTBI group there were significant differences between the left and the right Internal capsule at both time points after injury (mTBI, n=6; Sham, n=6; avg±SEM; Student t-test, *, p<0.05).

### DTI Results

FA maps are shown in **Figure 5** for both mTBI and Sham pigs.

**Figure 5.**
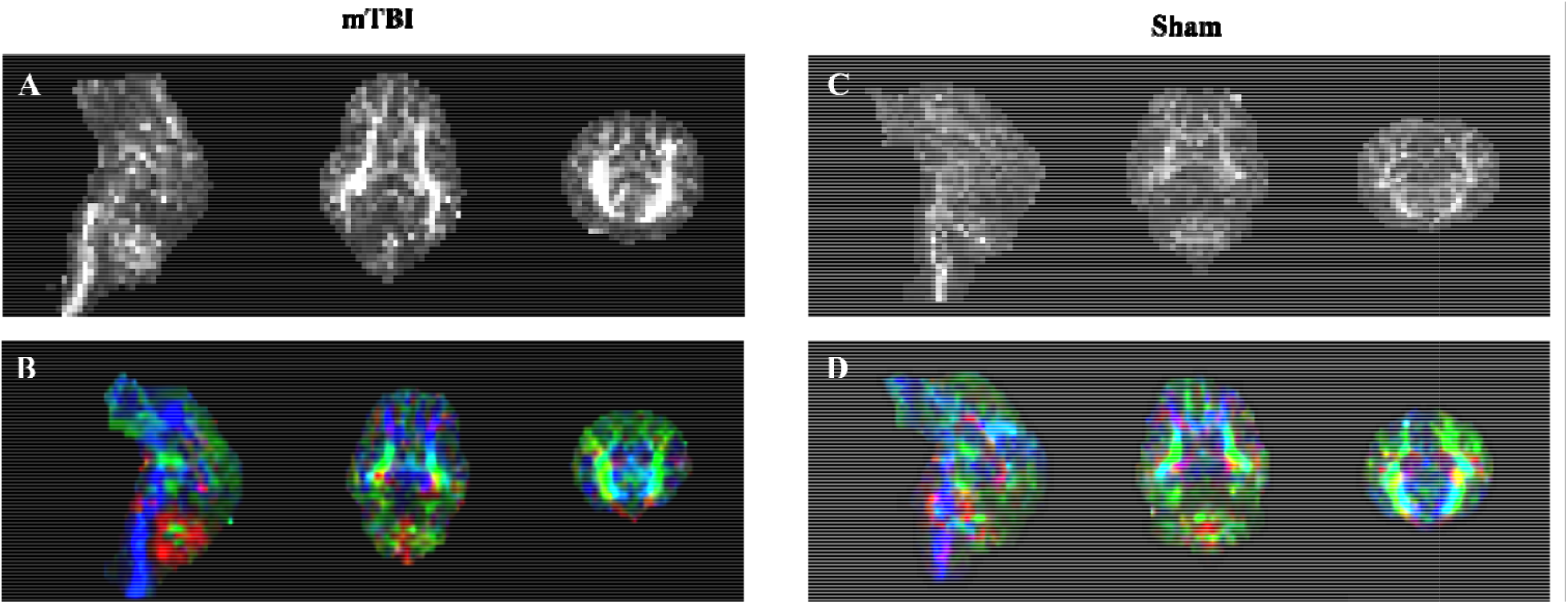
FA Maps in grayscale and color-coded orientations. Left: FA maps (A) in grayscale (sagittal, transverse, and coronal planes respectively), and FA maps (B) in color code for one mTBI pig. Right: FA maps (C) in grayscale, and FA map (D) in color code for a sham pig. Color codes for orientation are as follows: Red indicates left to right, Green represents anterior to posterior, and Blue denotes superior to inferior.

**Table 2** shows the DTI parameters of white matter integrity that were calculated for each subject at 1-month and three-month post-injury.

**Table 2.**
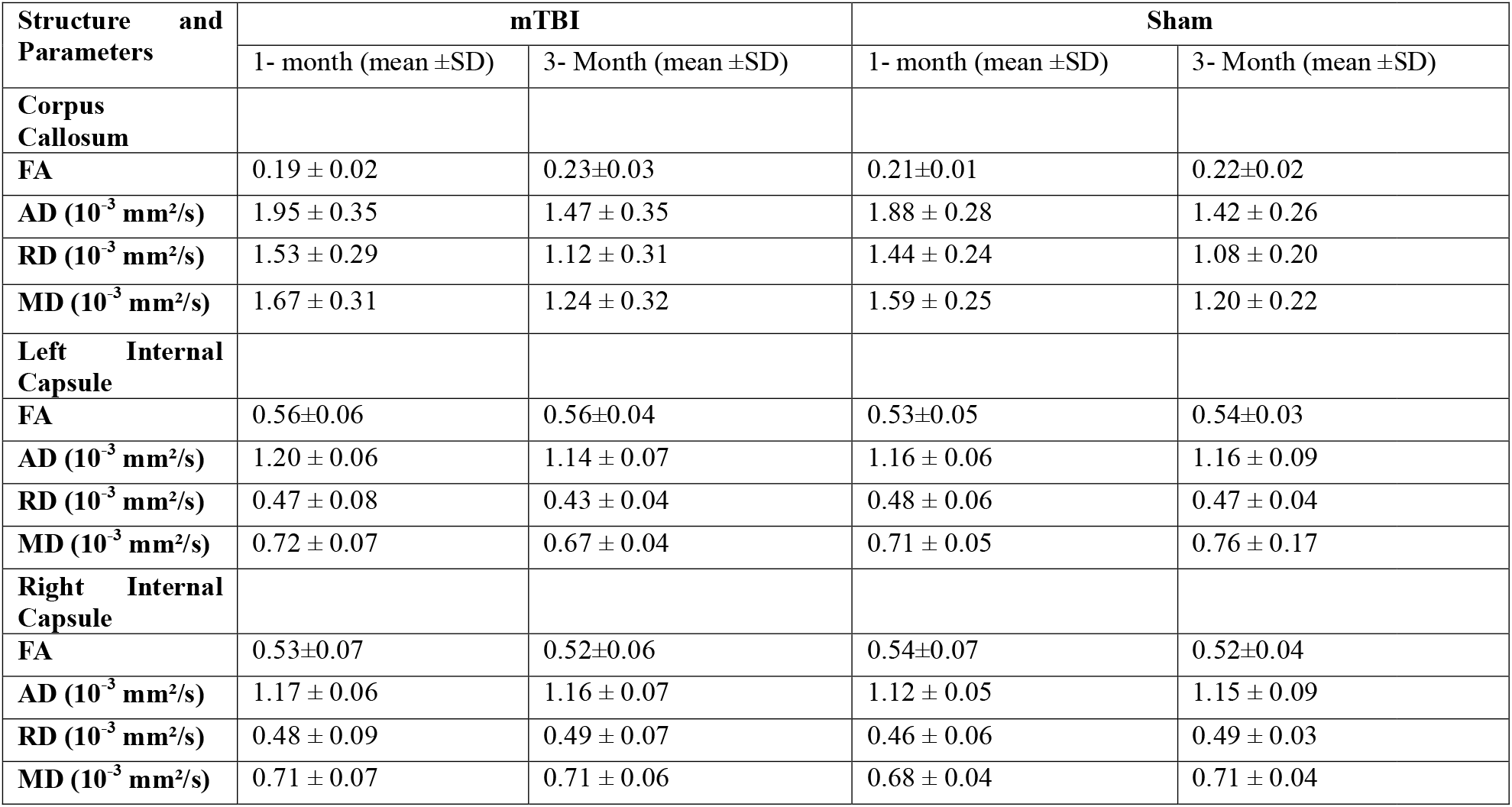
**DTI parameters of white matter tracts for the mTBI and Sham groups at 1-month and 3-months after injury** (mTBI, n=6; Sham, n=6).

| Structure and Parameters | mTBI |  | Sham |  |
| --- | --- | --- | --- | --- |
| | 1- month (mean ±SD) | 3- Month (mean ±SD) | 1- month (mean ±SD) | 3- Month (mean ±SD) |
| **Corpus Callosum** |  |  |  |  |
| FA | 0.19 ± 0.02 | 0.23±0.03 | 0.21±0.01 | 0.22±0.02 |
| AD ($10^{-3}$ mm<sup>2</sup>/s) | 1.95 ± 0.35 | 1.47 ± 0.35 | 1.88 ± 0.28 | 1.42 ± 0.26 |
| RD ($10^{-3}$ mm<sup>2</sup>/s) | 1.53 ± 0.29 | 1.12 ± 0.31 | 1.44 ± 0.24 | 1.08 ± 0.20 |
| MD ($10^{-3}$ mm<sup>2</sup>/s) | 1.67 ± 0.31 | 1.24 ± 0.32 | 1.59 ± 0.25 | 1.20 ± 0.22 |
| **Left Internal Capsule** |  |  |  |  |
| FA | 0.56±0.06 | 0.56±0.04 | 0.53±0.05 | 0.54±0.03 |
| AD ($10^{-3}$ mm<sup>2</sup>/s) | 1.20 ± 0.06 | 1.14 ± 0.07 | 1.16 ± 0.06 | 1.16 ± 0.09 |
| RD ($10^{-3}$ mm<sup>2</sup>/s) | 0.47 ± 0.08 | 0.43 ± 0.04 | 0.48 ± 0.06 | 0.47 ± 0.04 |
| MD ($10^{-3}$ mm<sup>2</sup>/s) | 0.72 ± 0.07 | 0.67 ± 0.04 | 0.71 ± 0.05 | 0.76 ± 0.17 |
| **Right Internal Capsule** |  |  |  |  |
| FA | 0.53±0.07 | 0.52±0.06 | 0.54±0.07 | 0.52±0.04 |
| AD ($10^{-3}$ mm<sup>2</sup>/s) | 1.17 ± 0.06 | 1.16 ± 0.07 | 1.12 ± 0.05 | 1.15 ± 0.09 |
| RD ($10^{-3}$ mm<sup>2</sup>/s) | 0.48 ± 0.09 | 0.49 ± 0.07 | 0.46 ± 0.06 | 0.49 ± 0.03 |
| MD ($10^{-3}$ mm<sup>2</sup>/s) | 0.71 ± 0.07 | 0.71 ± 0.06 | 0.68 ± 0.04 | 0.71 ± 0.04 |

Results indicate that mTBI pigs show a significant increase in FA values of the corpus callosum at 3 months compared to 1 month (paired t-test, p=0.037). Sham pigs did not show any difference in FA values over this time period. (**Figure 6A**). Analysis of MD, RD, and AD values demonstrated that there is a significant decrease between 1-month to 3-months post procedure, in both mTBI and Sham animals, suggesting that these variables reflect changes in brain maturation occurring across all groups.

**Figure 6.**
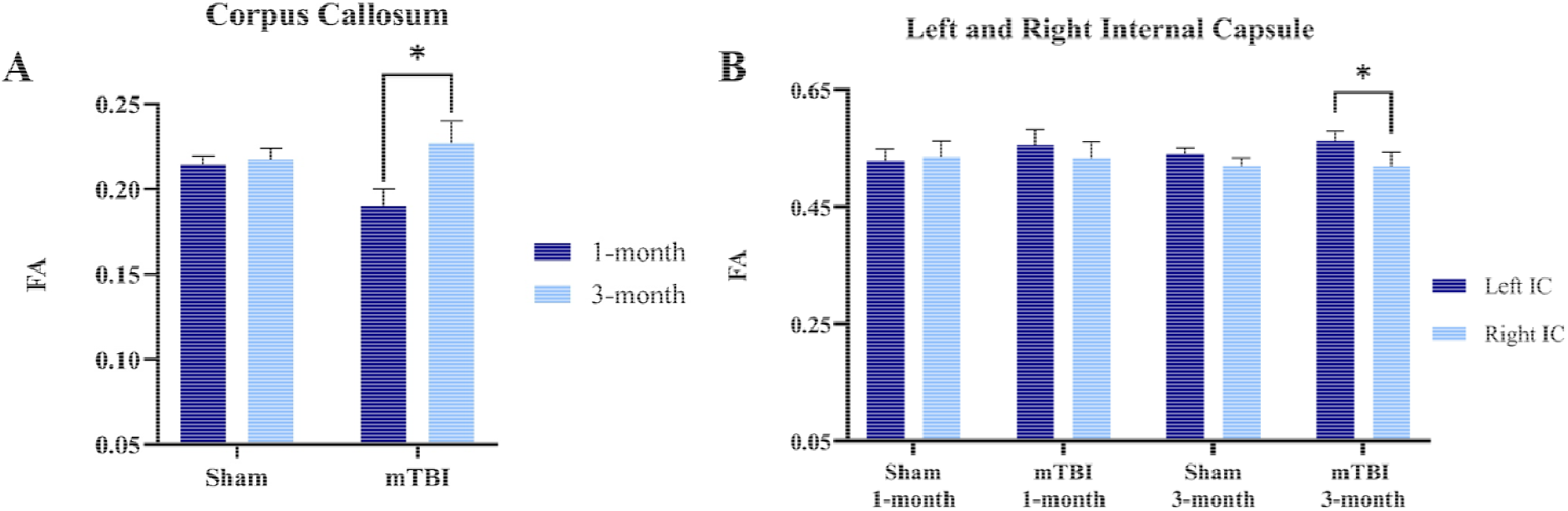
FA Changes in Corpus Callosum and Internal Capsule post-mTBI. A) FA changes in the Corpus Callosum between 1-month to 3-month post-injury. The graphs depict the mean differences between mTBI and Sham groups at the two time points. mTBI group shows a significant increase in FA from 1-month to 3-month post-injury. B) The comparison of FA values between Left and right Internal Capsule for both group and each time point. At 3-month post injury, the FA values in the right internal capsule are significantly lower compared to the left one in the mTBI group (mTBI, n=6; Sham, n=6; avg ±SEM; Student t-test, *, p<0.05).

The DTI variables were calculated for the right and left Internal capsule which is part of the cort cospinal pathway. The results show that there was a significant difference in FA values between the right and the left Internal capsule at 3 months after injury in mTBI pigs and is illustrated in **Figure 6B**. Specifically, the Right internal capsule showed lower FA values compared to the left one, which is consistent with the location of the TBI impact to be over the right frontal cortex.

Post-mortem histology of LFB, an indicator of white matter, has been performed. **Figure 7** shows representative LFB staining on 4 microns brain slice obtained from the frontal cortex. An area within the Corpus callosum was analyzed, and the results indicate that mTBI pigs showed a significant decrease of 3.4% in LFB staining compared to Sham (mTBI, n=3; Sham n=3; Student t-test, p<0.05).

**Figure 7.**
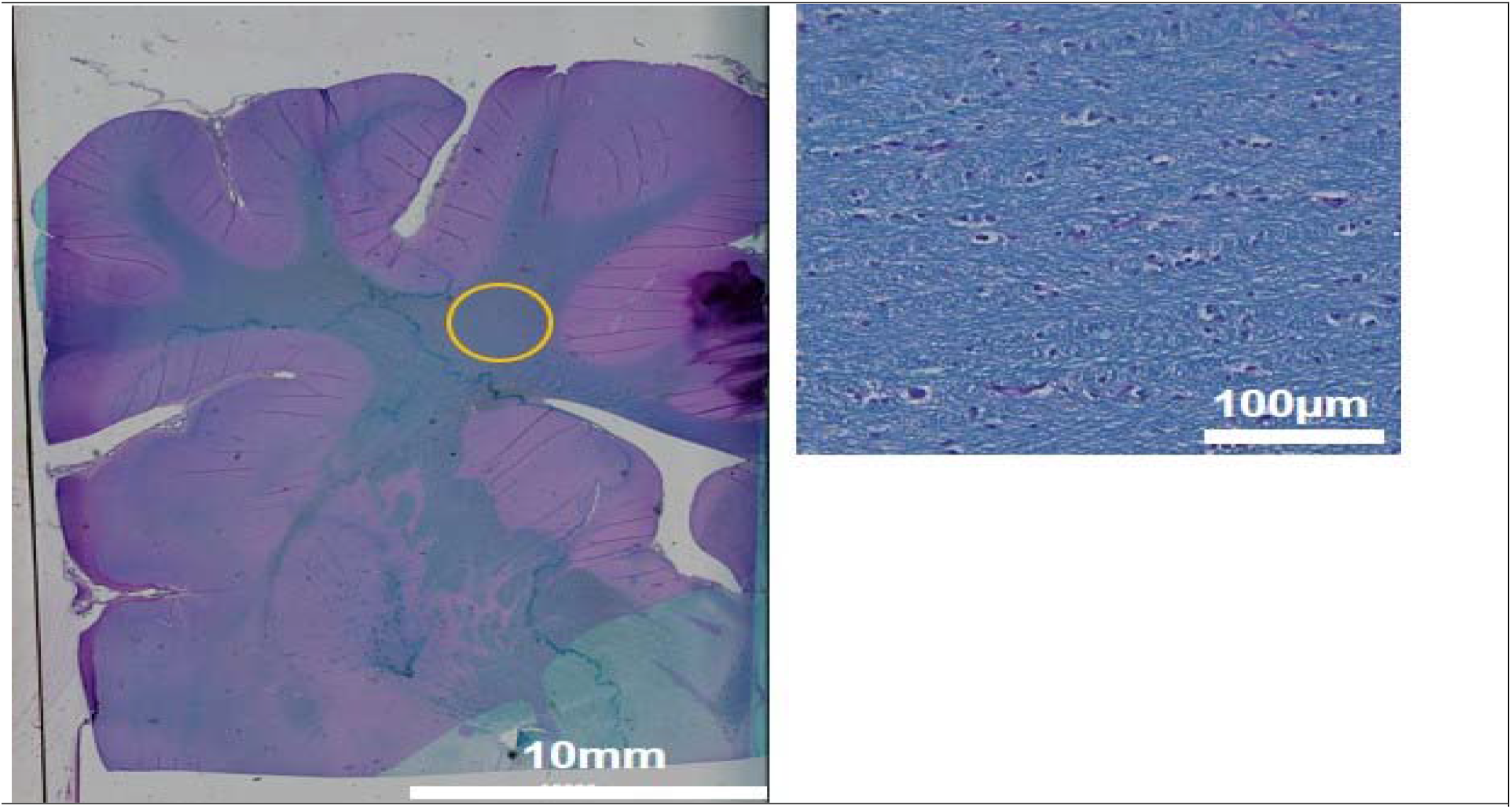
Histological analysis. Luxol fast blue histology. Left: An image of 4 µm thick brain slice stained for Luxol fast blue showing the area of the corpus callosum that was analyzed to evaluate myelin in the pig’s brain. Right: A 20x magnification of the corpus callosum area shows myelin architecture with high resolution.

## Discussion

The characteristics of the closed-head mTBI model in adolescent pigs mimicked an injury with a clinically relevant magnitude, comparable to mTBI in pig models reported by other groups,^52^ and coincides with the magnitude of injury reported in youth sports.^53-55^ The model induced long-term changes in the integrity of white matter tracts and brain structures that may be clinically translational. A longitudinal study that incorporates age-comparable reference data offers insights into the maturation of white matter over time which is important for assessing MRI data from children due to their developing brain.^56^ Our study’s use of a youth animal model with a unique image acquisition timeline minimizes age-related variability in the results.^57^

In this study, we concentrated on critical white matter tracts, specifically the internal capsule and corpus callosum, consistent with previous research.^58^ The internal capsule comprises axons connecting the cortex and thalamus. Both the left and right internal capsules contain muscle fibers that transmit information from the contralateral side of he body (excluding the upper face, which has bilateral connections). Consequently, damage to the internal capsule would result in paralysis or severe weakness on the entire opposite side of the body. Additionally, many axons traversing the upper brain stem are also part of the internal capsule. The corpus callosum is particularly susceptible to traumatic injury, regardless of biomechanical factors or injury severity.^59,60^ Rapid degeneration of unmyelinated fibers in the injured corpus callosum was observed.^61^

Previous research has documented a rise in FA values in the corpus callosum approximately three days post-injury in adults and six days post-injury in young children.^62^ Increased anisotropic diffusion can be more pronounced and prolonged in youth mTBI.^63^ A study that replicated their earlier results found higher anisotropic diffusion and lower radial diffusivity in the genu of the corpus callosum.^64^ The observed increase in FA values is likely due to axonal swelling, extended mild cytotoxic edema,^65^ neuroinflammation, occurring during the sub-acute phase of injury.^66^ Secondary cerebral swelling; axonal injury; and inflammation and regeneration also contribute to secondary injury in infants. In current research, FA in the corpus callosum for the mTBI pigs increased over time from 1-month to 3-months indicating callosal damage.^67^ While an increase in FA might suggest a normal myelination process over time,^68,69^ the changes observed in the sham group imply that the FA alterations may not be attributable to typical axonal development in the youth brain. Studies have detected unusually high FA in the corpus callosum^70^ and internal capsule^58^, which is thought to result from an influx of intracellular water or changes in the concentration of extracellular water restricting the diffusion^71^ more frequently observed in a developing brain. Another study has evaluated it as reorganization of WM tract due to plasticity^72^ An upward trend in fractional anisotropy (FA) is frequently observed in athletes or individuals with sports-related concussions. This trend is hypothesized to result from processes such as axonal budding^73^ and growth, or the formation of glial scar.^74^ Research utilizing Neurite Orientation Dispersion and Density Imaging (NODDI) parameters further supports the hypothesis that there is an increase in neurite density following injury.^75^

Within the mTBI groups, changes in internal capsules were not significant over time, unlike in the corpus callosum. However, a temporal effect was observed, as FA decreased over time in the right internal capsule compared to the left in the mTBI group indicative of injured white matter area. The reduction in FA values in the right internal capsule compared to the left at 3 months post injury is likely attributable to the impact affecting the ipsilateral side of the internal capsule significantly. Such reduction in FA during late post injury may reflect disruption in axonal transport, axonal swelling, demyelination or Wallerian degeneration. Existing literature on youth mTBI model showed lower FA in multiple regions but no changes in Corpus Callosum.^76^ Results showed significant differences in diffusion characteristics between affected white matter regions and contralateral homologous regions (Left and Right Internal Capsule) in patients with mTBI.^77^ The current findings also indicate that the decrease in FA over the 3 months after the injury could be indicative of Wallerian degeneration in the affected neurons.

Different trends in diffusion changes associated with mTBI have been documented; In human youth mTBI model both higher^78^ and lower FA^79^ were found. In the present study, we did not detect a group effect. Indeed, previous research has shown minimal differences between mTBI and sham groups, suggesting that the TBI group was only mildly affected, with subtle changes in white matter structure.^27^ Unlike our findings, other ROI-based studies have identified significant group differences in several brain areas associated with mild traumatic brain injury (TBI). Notable differences were observed in the genu and splenium of the corpus callosum and the posterior limb of the internal capsule.^79^ Additionally, differences were found in the anterior corona radiata, uncinate fasciculus, and cortico-spinal tract, as well as in the sagittal stratum and superior longitudinal fasciculus.^80-82^ However, null findings have also been reported, revealing no differences between groups in 20 white matter tracts,^83^ corpus callosum,^81^ VBM analysis^84^ and TBSS study on athletic concussion.^85^ Therefore, the current findings offer evidence of diffusion abnormalities.

Another study on closed head injury animal models emphasized the dominance of cellular edema during acute period of post-injury.^86^ A Case study of an infant observed a transient increase in both FA and MD in gray matter regions reflecting complex interaction between cytotoxic and vasogenic edema.^87^ Sometimes the higher FA is believed to be due to restricting the sample to voxels with more homogeneous fiber orientation.^88^ Another study examining the longitudinal changes in the corpus callosum microstructure in children found a significant increase in FA only in the sham group, supporting the notion of abnormal white matter maturation in children with mTBI.^67^

In examining brain volume alterations in mTBI, regions like the corpus callosum, thalamus, hippocampus, hypothalamus, and cerebellum are regarded as particularly susceptible due to their extensive connections with other brain areas. Studies specifically focusing on hippocampal volume are scarce, but any observed changes in this region could be significant for understanding the long-term impacts of mTBI.^89^ For the thalamus, while most research indicates atrophy, a study by Munivenkatappa et al.^90^ reported an increase in thalamic volume at a three-month follow-up. Similarly, Obermann et al.,^91^ suggested that increases in gray matter in the thalamus and cerebellum might result from adjustments in pain tolerance following injury. Conversely, although general brain atrophy is commonly observed, some studies have found abnormal enlargement in subcortical regions like the thalamus, amygdala, and hippocampus, potentially due to hyperactivity, hypertrophy, and neuroinflammation.^92^

The present study did not identify any significant group-level differences in white or gray matter volumes related to mTBI but showed longitudinal in-group changes. Previous research has reported changes in white matter volume in youth patients from post-acute scans, but no changes in gray matter volume.^18^ Volumetric analysis shows a significant increase in brain volume for the corpus callosum only in the mTBI group supports the theory that traumatic brain edema causes swelling and increases brain tissue water content.^93,94^ The white matter areas such as the left and right internal capsules, mTBI pigs showed more significant volume changes over time compared to Sham. Additionally, only mTBI pigs exhibited a significant expansion in volume in gray matter regions, including the cortex, thalamus, and hippocampus, likely due to brain edema. Quantitative analyses have shown significant atrophy in gray matter regions such as the cerebral cortex, thalamus, and hippocampus within 2 months to 1-year post-injury in rats.^37,95^ A longitudinal study found that traumatic brain injury (TBI) leads to ongoing atrophy in the corpus callosum, thalamus, and hippocampus for up to 1 to 5 years, with an increase in ventricular volume, indicating a prolonged impact on brain volumes.^96^ However, another study reported no significant volume loss in these gray matter regions, as well as in white matter regions like the internal capsule and corpus callosum. Notably, all studies observed significant expansion of the lateral ventricles.^97^ In both analyses, the impact has clearly impeded the normal maturation of white matter. The mixed model analysis indicates that the interaction between gender and group is not significant (p > 0.05) for both the Corpus Callosum and the Left and Right Internal Capsule, suggesting that gender does not have an effect on the results.

One constraint of our study pertains to the modest sample size employed. Despite our efforts to examine correlations among mean FA values during the subacute and early chronic phases in specific regions exhibiting notable changes over time, no significant correlations were identified. However, the absence of statistical significance may be attributed to the limited sample size utilized. Future studies with increased sample size may reveal DTI changes that have been too subtle to be identified here. The use of representative regions rather than the whole white matter structures may have introduced some bias into the results. Furthermore, ROI analyses can be problematic due to the low resolution of DTI maps and the unavoidable partial volume effects. The corpus callosum is especially susceptible to partial volume effect due to adjacent cerebrospinal fluid, with this effect being more pronounced when DTI voxels have bigger size along the z-axis.^98^ Thus, this method may diminish sensitivity to small lesions. It will be important to obtain pre-injury DTI metrics to identify intra-individual changes. The use of varying approaches can occasionally lead to inconsistencies in the results. This highlights the sensitivity of DTI to white matter damage during both the sub-acute and chronic phases, underscoring the importance of conducting longitudinal studies or follow-up assessments post-injury. A study found that in children with mTBI, abnormal white matter maturation in the corpus callosum was resolved within 6 to 8 months, suggesting the need for more research into white matter recovery and the developing brain’s vulnerability to mTBI.^67^

In conclusion, this work shows MRI biomarkers and histopathology evidence of white matter injury in a miniature pig model of mTBI. DTI and volumetric analysis of MRI data demonstrate that mTBI results in significant changes in white matter integrity. This could be important both for diagnostic purposes and for clinical management of patients.

### Transparency, Rigor and Reproducibility Summary

Rigor and reproducibility: The design and statistical analysis associated with this project was performed with a Biostatistician. P-values were corrected by multiple testing using Bonferroni correction. Basic statistical inference was performed in all points of measurements. To determine statistical significance between mean of groups, we used t-test (for experiments with two categories). mTBI sequela effects was tested by gender as a variable. The statistics was done with R and GraphPad prism. Randomization: Pigs were randomly assigned to experimental and sham groups. Both genders were included. Blinding: Each cohort of pigs included both experimental and sham animals. The veterinary staff who performed the surgical procedures kept the health records for each individual and assigned a code. Apart from the injury itself, both experimental and sham-control pigs underwent similar surgical procedures and received similar post-operative treatment. This eliminated the ability for blinded observers to easily tell which pig has received an impact. The investigators who perform the MRI imaging analysis received only the code for each subject and were blinded to the group it belongs to. The power analysis was based on preliminary data consisting of mTBI and control groups. The preliminary data showed that the interindividual variation is higher than the residual variance and a sample size of n=6 experimental pigs, and n=6 control pigs is needed to reach a power of 0.8 and a significance level of 0.05 (2-sides).

